# Using the DNA methylation profile of the stress driver gene *FKBP5* for chronic pain diagnosis

**DOI:** 10.1101/2022.12.22.521573

**Authors:** Maria Maiarù, Richard J. Acton, Eva L. Woźniak, Charles A. Mein, Christopher G. Bell, Sandrine M. Géranton

## Abstract

Epigenetic changes can bring insight into gene regulatory mechanisms associated with disease pathogenicity, including chronicity and increased vulnerability. To date, we are yet to identify genes sensitive to epigenetic regulation that contribute to the maintenance of chronic pain and with an epigenetic landscape indicative of the susceptibility to persistent pain. Such genes would provide a novel opportunity for better pain management, as their epigenetic profile could be targeted for the treatment of chronic pain or used as an indication of vulnerability for prevention strategies. Here, we investigated the epigenetic profile of the gene *FKBP5* for this potential, using targeted bisulphite sequencing in rodent pre-clinical models of chronic and latent hypersensitive states. The *FKBP5* promoter DNA methylation (DNAm) signature in the CNS was significantly different between models of persistent pain and there was a significant correlation between CNS and peripheral blood *FKBP5* DNAm, indicating that further exploration of *FKBP5* promoter DNAm as a biomarker of chronic pain pathogenic origin is warranted. We also found that maternal separation, which promotes the persistency of inflammatory pain in adulthood, was accompanied by long-lasting reduction in *FKBP5* DNAm, suggesting that *FKPB5* DNAm profile may indicate the increased vulnerability to chronic pain in individuals exposed to trauma in early life. Overall, our data demonstrate that the *FKBP5* promoter DNAm landscape brings novel insight into the differing pathogenic origins of chronic pain, may be able to diagnose and stratify patients, and predict the susceptibility to chronic pain.

## Introduction

Chronic pain is a significant burden to society. In the UK alone, it affects 13-50% of adults, with at least 10% of those living with moderate to severe disabling pain [5,31]. Importantly, chronic pain is not only limited to older age groups and affects up to 30% of those aged 18 to 39 years [16,31]. Unfortunately, managing this condition remains a major challenge for clinical practice. Failure to diagnose and classify chronic pain patients may certainly contribute to the lack of efficacy of present therapies that are also often accompanied by significant side-effects, including the potential for addiction [7,39]. An opportunity for better pain management lies with strategies preventing the transition from acute to chronic pain. Unfortunately, identifying vulnerable individuals and preventing their pain to persist remains challenging [13,30]. Thus, understanding more precisely the mechanisms involved in the pathogenesis of chronic pain may uncover novel therapeutic avenues as well as opportunities for more targeted personalised medicine.

The protein FKBP51, encoded by the gene *FKBP5*, is a regulator of the stress axis [12,17]. Its genetic deletion and pharmacological blockade alleviate persistent pain states in rodents [27,28,40], while identified genetic variants have been associated with pain sensitivity after trauma in early small scale human studies [1,4,37]. However, these genetic findings have not, to date, been replicated in larger biobank-scale GWAS. Importantly, persistent pain driven by FKBP51 is associated with a rapid decrease in *FKBP5* DNA methylation (DNAm) in the rodent spinal cord which correlates with the spinal increase of the FKBP51 protein that can be locally prevented to relieve established persistent pain [27,28]. Whether this change in DNAm is maintained together with the hypersensitive state has not explored but compelling evidence for a role of *FKBP5* DNAm dysregulation in human chronic diseases has recently been provided through genome wide DNA-methylation studies [3,8,18,34,44].

Especially relevant in the context of preventive approaches for the management of chronic pain, early work from Klengel et al. indicated the potential that early-life trauma in humans may lead to a decrease in *FKBP5* DNAm, an epigenetic change that primes *FKBP5* for hyperresponsiveness and may increase the susceptibility to post-traumatic stress disorder (PTSD) in adulthood [20]. We hypothesise that similar processes could underlie the vulnerability to chronic pain [14], but we do not know whether stress exposure could induce a change in DNAm at spinal cord level. Such a reduction in DNAm could prime *FKBP5* for hyper-responsiveness to injury in later life, therefore increasing the likelihood of developing persistent pain. This would provide a mechanistic basis for the well-described link between early life trauma and increased susceptibility to chronic pain [6].

The hypothesis of this project was that significant trauma of physical or emotional nature could induce a long lasting change in the DNAm profile of the *FKBP5* gene in the central nervous system (CNS), which could maintain an individual in a hypersensitive state, or leave an individual susceptible to chronic pain. This was investigated through the use of pre-clinical models of persistent pain and of latent states of hypersensitivity.

## Methods

### Animals

Subjects in all experiments were male Sprague-Dawley rats obtained from the colony at UCL. All rats were kept in their home cage in a temperature-controlled (20° ± 1° C) environment, with a light-dark cycle of 12 hours. Home cages were enriched with tubes, nesting materials and chewing sticks. Food and water were provided ad libitum. All behavioural experiments were performed by the same experienced female experimenter, who was always blind to treatment group for all behavioural tests. Animals were allocated to a treatment group using a random number generator. Mechanical hypersensitivity was measured using a series of calibrated Von Frey’s monofilaments [28]. The threshold was determined by using the up-down method [9]. The data are expressed as log of mean of the 50% pain threshold ± SEM.

### Animal models

#### Ankle joint inflammation

We induced joint inflammation with the injection of Complete Freund’s Adjuvant (CFA; 10 μl) in the ankle joint in adult male rats (250g, 7-week old) as before [15]. Sham animals were only put under anaesthesia for few minutes. For molecular analysis, tissues were collected 30 days after the CFA injection.

#### Spared Nerve Injury

For the neuropathic pain model, we used spared nerve injury (SNI) in adults male rats (250g, 7-week old) as before [10,28]. Sham animals were put under anaesthesia and the nerve was exposed but untouched before closing the wound. For molecular analysis, tissues were collected 30 days after the SNI surgery.

#### Physical priming with inflammation

The mechanical hypersensitivity that develops after plantar subcutaneous (s.c.) injection of carrageenan resolves within few hours in neonates at Postnatal day 5 (P5). However, when these animals are re-injured later on in life, they display a greater mechanical hypersensitivity when compared with animals that have not previously been injected with carrageenan [25]. Primed male rats received carrageenan (1ul/g of body weight) at P5 in the left hindpaw and control animals were only put under anaesthesia for few minutes. All male rats received a plantar s.c. injection of PGE2 in the left hindpaw (100ng/25ul) at P40. For molecular analysis, tissues were collected at P40, with no injection of PGE2.

#### Emotional priming with early life adversity: maternal separation

Stress in early life was induced in male rats by maternal separation, 3h/day from P2 to P12 [19]. Control male rats stayed with the dam in their home cage. All experimental male rats received a plantar s.c. injection of PGE2 (100ng/25ul) at P42. For molecular analysis, tissues were collected at P40, with no PGE2 injection.

#### DNA & RNA extraction

Following behavioural assessment, animals were terminally anesthetized with CO_2_. CNS samples (full hippocampus and ipsilateral quadrant of the L4-L6 spinal cord segment) and blood were harvested from animals and DNA and RNA was extracted from these same samples. We used the All Prep DNA/RNA/miRNA Universal Kit from Qiagen, https://www.qiagen.com/us/products/discovery-and-translational-research/dna-rna-purification/multianalyte-and-virus/allprep-dnarnamirna-universal-kit/?catno=80224 as per the manufacturer’s instructions.

#### RTqPCR

Reactions were performed at least in triplicate, and the specificity of the products was determined by melting curve analysis. The ratio of the relative expression of target genes to ⍰-actin was calculated by using the 2^⍰Ct^ formula as before [28].

#### DNA methylation Targeted Bisulphite Sequencing

Primers for targeted DNA methylation (DNAm) bisulphite sequencing were designed with MethPrimer 1.0 [24] to survey of the status of the potentially functional CpG dense promoter region of the *FKBP5* gene. This includes the 3 upstream CpG islands (CGIs) and their shore regions. (Supplementary Figure S1). Parameters employed for the primer design were Product Size (minimum 125bp; optimal 150bp; maximum 175bp), Primer Tm (min. 50°; opt. 55°; max. 60°), Primer Size (min 20bp; opt. 25bp; max 30bp), and including at least 4 product CpGs. 21 loci were initially targeted covering 90 CpGs (Supplementary Table S1).

The targeted sequencing loci were assessed for DNAm changes between the various animal models. Of all the targeted loci, reliable results passing QC were generated for 90 CpGs. Quality control included visualisation assessment pre- and post-read trimming via fastqc v0.11.5 (Andrews, S. FastQC, Babraham Bioinformatics, Cambridge, 2010) and multiqc v1.0[11]. Cutadapt v1.13 [29] and a custom perl5 script was employed to trim targeting primers, with then quality trimming via trim_galore 0.5.0 [22]. Bismark v0.20.0 [21], incorporating the bowtie2 v.3.1 [23] aligner, was used for DNAm calling. Alignment was performed against targeted loci ±100 bp in the rat rn6 genome with a minimum coverage threshold of 25 reads. Differential methylation analysis was performed using RnBeads v2.0.1 [32].

#### Cell-type heterogeneity correction and reported differentially methylated cytosines

To account for influence of the cell-type heterogeneity of the DNA samples, RTqPCR data was used to correct CNS samples for cell type proportions. One correction was applied using as covariates Aif1 for glia, Aldhl1 for astrocytes and Olig1 for oligodendrocytes (Cell Type Correction 1, CTC1). To check the robustness of our approach, another correction was also applied using as covariates CxCr1 for microglia, Aldhl1 for astrocytes and Olig1 for oligodendrocytes (Cell Type Correction 2, or CTC2). CpGs for differentially methylated loci were tested with Wald test as implemented in RnBeads [32].

Our final high quality CpG results reported here were required to pass a threshold of (1) coverage greater than 300 reads for all samples within a group; (2) methylation difference greater than 1% and (3) P<0.05 for both CTC1 and CTC2 (or P<0.1 for the reported trend results). In the Results section, the P values for CTC1 are stated. We were unable to correct the blood samples for cell type variation due to a lack of haematological data, however, these were considered as pilot data to explore the potential biomarker utility of blood cell-derived DNAm.

#### Transcription Factor Binding Motif Enrichment

The Transcription Factor Affinity Prediction (TRAP) v3.0.5 [35] ‘single sequence’ option was used to explore any enrichment for Transcription Factor (TF) binding motifs surrounding significant CpGs. The sequences around each CpG were expanded to +/-10bp and the FASTA sequence extracted (rn6) via UCSC. The Transfac 2010.1 Vertebrate matrix set was interrogated with mouse_promoter set as background model.

## Results

### CFA-induced ankle joint inflammation and nerve injury have different FKBP5 promoter DNA methylation landscape

Injection of CFA (10μl) in the ankle joint and SNI surgery induced long-lasting mechanical hypersensitivity in adult male rats (Fig.1A; Day3 to Day30: CFA vs CFA Sham: F_1,15_=84, p<0.0001; SNI vs SNI Sham: F_1,11_=48; p<0.0001). There was no difference in the degree of mechanical hypersensitivity induced by the two models. We had previously found that both CFA injection in the ankle joint and SNI induced an up-regulation of *FKBP5* expression in the dorsal horn within few hours of the initiation of the pain states [27,28]. Here, using RTqPCR to measure *FKBP5* mRNA levels in the maintenance stage of the pain states, we found no change in expression of *FKBP5* in the hippocampus or in the dorsal horn following either CFA (30 days post-CFA; Fig.1B) or SNI (30 days post-surgery; Fig.1C) when compared to their respective sham groups.

**Figure 1:**
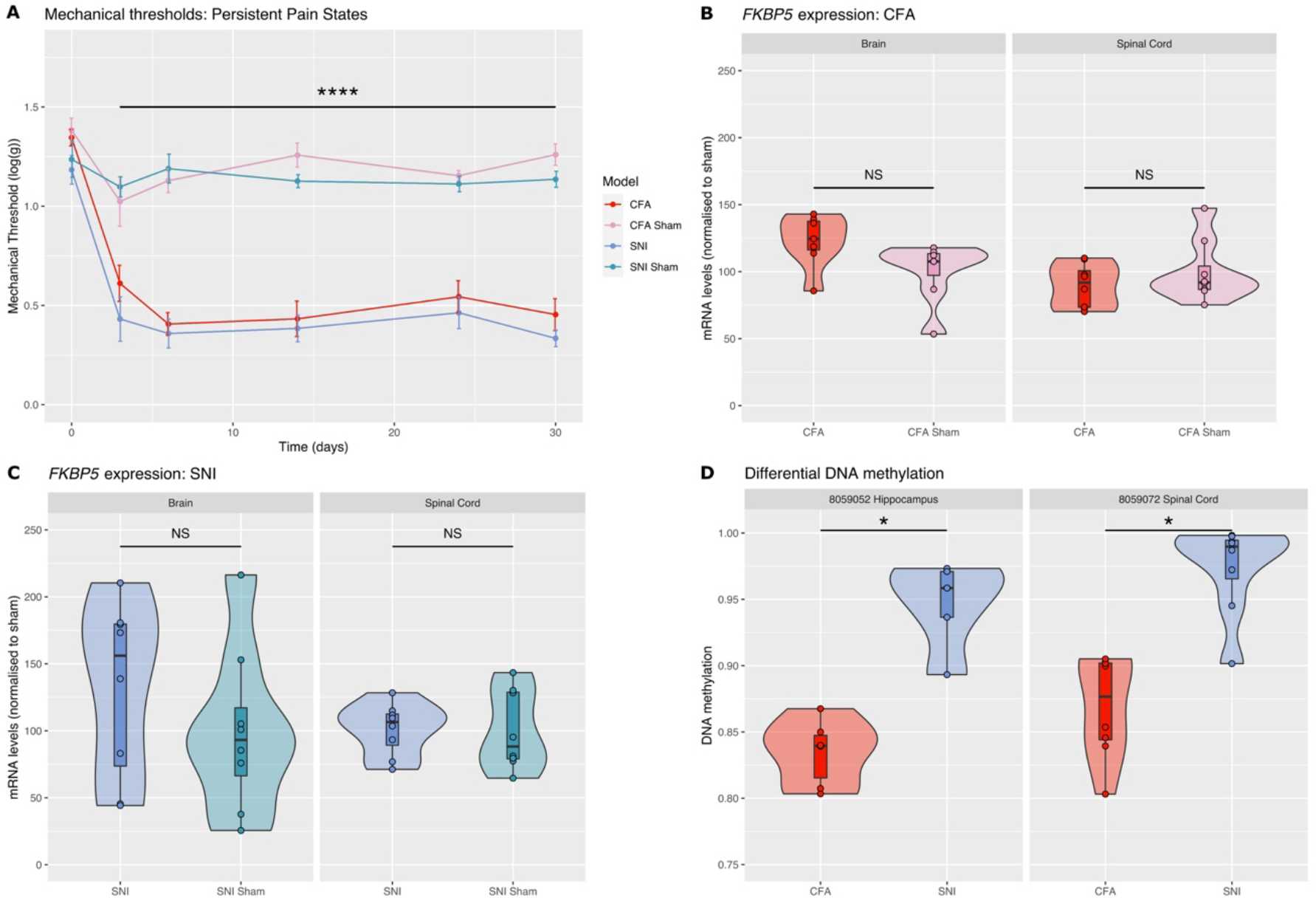
CFA-induced ankle joint inflammation and nerve injury induce mechanical hypersensitivity of similar degree, no long-lasting changes in FKBP51 expression but differences in FKBP5 promoter DNA methylation. **(A)** Adult male rats were injected with CFA (10μl) in the ankle joint or had SNI surgery on day 0 (N=8/8/8/8). Both models induced significant mechanical hypersensitivity compared with their sham groups, but there was no difference in the mechanical hypersensitivity between the two models. (****P<0.0001, CFA vs CFA Sham and SNI vs SNI Sham). (**B-C)** There was no significant change in FKBP51 mRNA expression in the spinal cord and the hippocampus 30 days following CFA and 30 days following SNI. mRNA levels were quantified by RTqPCR (N=8/8). Data was normalized to Sham average. (**D)** We identified two CpG loci (8,059,052 & 8,059,072) with different levels of methylation between CFA and SNI tissue, in the Hippocampus and in the spinal cord, respectively. (*P<0.05).

However, we identified one CpG with greater levels of methylation in the SNI animals compared with CFA, in the hippocampus: CpG at positions chr20:8,059,052 (DNAmΔ = 11.2%; FDR-corrected p = 0.018) and one CpG with greater levels of methylation in SNI compared with CFA in the spinal cord: chr20:8,059,072 (DNAmΔ = 10.5%; FDR-corrected p = 0.023)(Fig.1D). While there was no significant difference in DNAm between the CFA and CFA Sham animals at these sites, there was a trend for an increase between SNI and SNI Sham at each CpG site (DNAm Δ = 3.2%; at positions chr20:8,059,052 and DNAmΔ = 8.5% at chr20:8,059,072, Supplementary Table S2).

We then interrogated the DNA sequence that the two CpGs (chr20:8,059,052 and chr20:8,059,072) reside centrally within for potential Transcription Factor Binding Site (TFBS) motifs. These two CpGs reside just beyond defined CpG shore regions (≤2kb from CGI) within shelf regions (>2kb and ≤4kb from the CGI, see Table 1). Including ±10bp of sequence around the probed cytosines, we explored the Transfac 2010.1 Vertebrate database for potential TFBS motifs via the Transcription Factor Affinity Prediction (TRAP) v3.0.5 tool ([35], see Methods). This identified three potential motifs for the two CpGs (see Supplementary Table S3). This included motifs for the TFs (1) E2F1 (E2F transcription factor 1), which was found elevated in the blood of patients with spinal cord injury induced neuropathic pain [45] and (2) AP1, recently identified as a key gene associated with neuropathic pain in the spinal cord [26].

**Table 1:** Location of significant DNA methylation difference between SNI and CFA tissue.

| Significant Cytosine (rn6) | Location | Distance to CGI (bp) | Nearest CGI | TFBS motifs ($\pm 10$bp) Transfac include | Mouse Syntenic Loci Regulatory Prediction (CNS) |
| --- | --- | --- | --- | --- | --- |
| chr20:8,059,052 | Intronic CGI shelf | 2,278 | chr20:8,061,330 -8,062,154 | E2F1 | Poised Enhancer / Quiescent Gene |
| chr20:8,059,072 | Intronic CGI shelf | 2,258 | chr20:8,061,330 -8,062,154 | AP1 | Poised Enhancer / Quiescent Gene |

Includes (Left to Right): Location of Cytosine (rn6 build); Location annotation; Distance to nearest CpG Island (CGI); Co-ordinates of nearest CGI; Highlighted enriched Transcription Factor Binding Site motifs in $\pm 10$ bp window centred on CpG via Transfac (TRAP analysis (24)); and observed Chromatin Segmentation signature in the syntenic Mouse location (GRCm38/mm10, via LiftOver) in developmental central neural tissue (27)).

Next, we looked for epigenomic evidence of regulatory activity at these CpG loci. Unfortunately, no Rat-specific Chromatin Segmentation currently exists, so we lifted the ±10bp sequences over to mouse genome (via UCSC LiftOver) where Chromatin Segmentation has been performed in developing mouse tissue (Encode Phase III)([38]). This revealed that these two CpGs resided within potential poised or primed Enhancer as well as Quiescent gene signatures in various developing neural tissues (Supplementary Figure S2).

### There is a correlation between the DNA methylation levels at chr20:8,059,052 in the hippocampus, the spinal cord and the blood

We observed that, while statistically not significant, the CpG at position chr20:8,059,052 was the second CpG - after chr20:8,059,072-most likely to show a change in DNAm in spinal cord tissue (DNAmΔ = 11.9%; FDR-corrected p = 0.1). Furthermore, while we were not able to statistically adjust and explore potential cell-type related DNAm variations in blood due to the lack of haematological cell-type data, we found an elevated level in DNAm at chr20:8,059,052 after SNI compared with CFA in blood of the same amplitude than that seen in the spinal cord and in the hippocampus (DNAmΔ = 8.5%; Bonferroni corrected p = 0.046) (see Supplementary Table S2). Finally, there was a significant correlation between the degree of DNAm in the blood and the degree of methylation in the hippocampus (Fig.2A), as well as between the degree of DNAm in the spinal cord and the degree of methylation in the hippocampus (Fig.2B).

**Figure 2:**
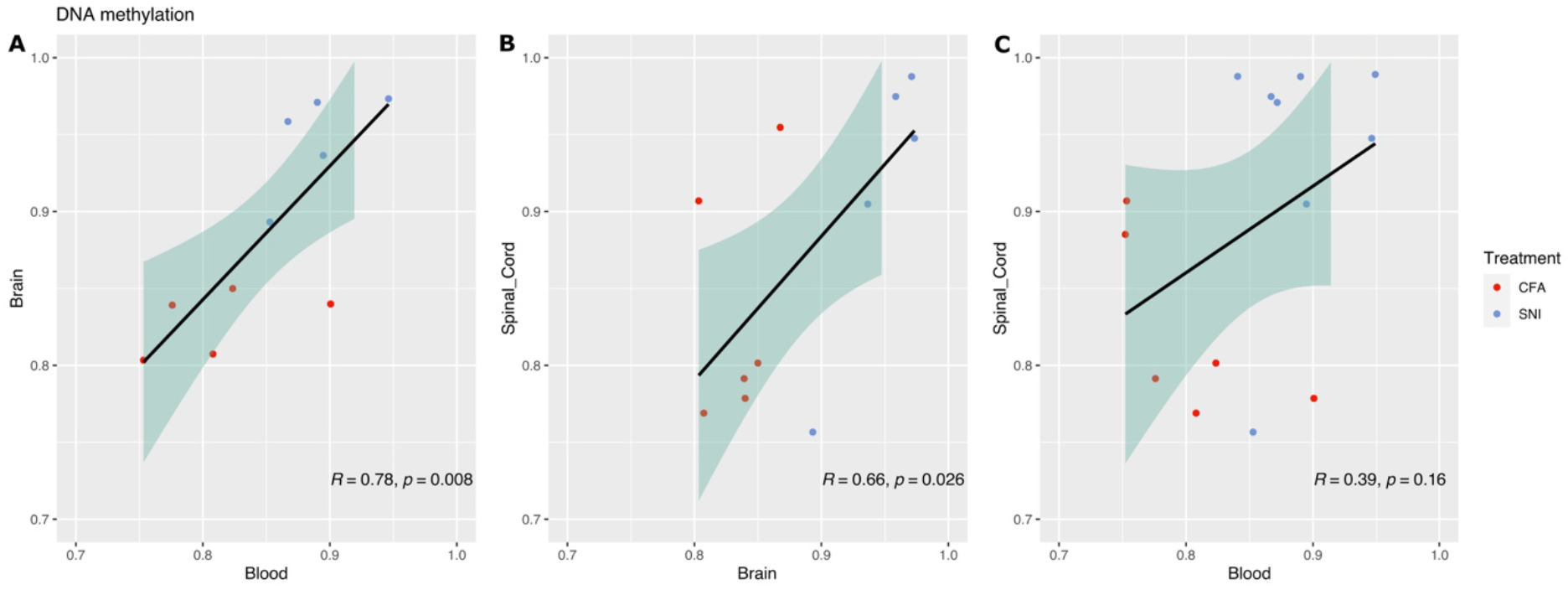
DNAm at chr20:8,059,052 in the hippocampus correlates with DNAm in the blood and in the spinal cord. **(A-B)** There is a significant correlation between DNAm in the hippocampus and the blood: Pearson’s correlation P=0.008, and between the hippocampus and the spinal cord: Pearson’s correlation: P=0.026. **(C)** The correlation between DNAm in the blood and the spinal cord did not reach significance.

### Short-lasting inflammation in early life primes for hyper-responsiveness to inflammation in adulthood but does not lead to a change in FKBP51 expression or FKBP5 DNA methylation in adulthood

Here, we used a model of early life inflammation which consisted in the injection of carrageenan (1μl/g of body weight) into the hindpaw of male rats at P5. This induced short-lasting mechanical hypersensitivity and primed the animals for hyper-responsiveness to injury in later life. This was indicated by the increased response to PGE2 seen in adult primed animals compared with adult control animals (Fig.3A). We then looked at *FKBP5* expression and DNAm in control animals and animals that had received an injection of carrageenan only, *i.e*. before the injection of PGE2, during the state of latent hyper-responsiveness. We found no change in *FKBP5* gene expression (Fig.3B) or *FKBP5* DNAm (data not shown).

**Figure 3:**
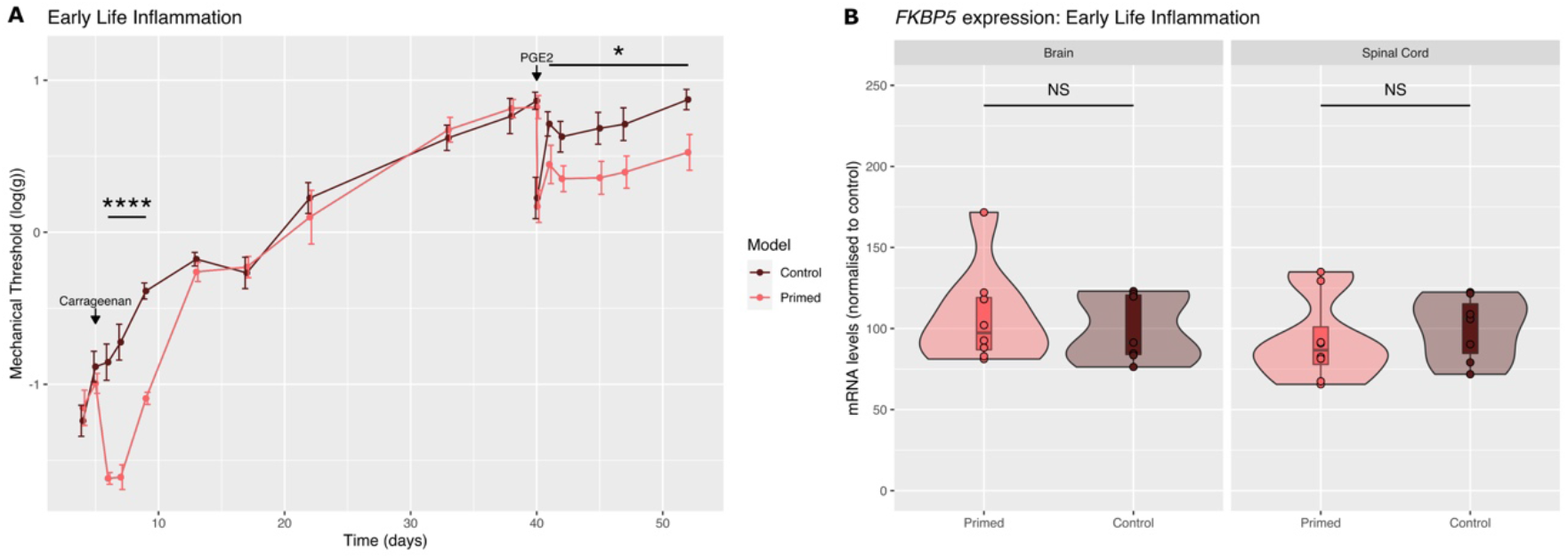
Intra-plantar inflammation in early life primes for hyper-responsiveness in later life but is not associated with changes in FKPB51 expression in the latent state of hypersensitivity. **(A)** Primed animals received s.c. plantar carrageenan (1μl/g of body weight) at P5 and all animals received an injection of PGE2 (100ng/25ul) at P38. N=8/8. 2-way ANOVA factor treatment, Day 5 to Day 9, F_1,14_=71.9, P<0.0001; Day 41 to Day 52, F_1,14_=7.3, P<0.05. Control vs Primed. (**B)** There was no change in FKBP51 mRNA expression in the spinal cord or the hippocampus 39 days after carrageenan injection. mRNA levels were quantified by RTqPCR. N=8/8. Data was normalized to Control.

### Early life adversity in the form of maternal separation primes for hyper-responsiveness to inflammation and leads to a decrease in FKBP51 expression and in FKBP5 DNA methylation in the spinal cord in adulthood

Finally, we explored the impact of early life adversity (Maternal Separation; Mat Sep) on inflammation-induced hypersensitivity, *FKBP5* expression and *FKBP5* DNAm in adulthood. Maternal Separation led to increased and prolonged mechanical sensitivity to PGE2 in adulthood (Fig.4A), as seen with early life inflammation. We then looked at *FKBP5* expression and DNAm in control animals and animals that had experienced maternal separation only, *i.e*. before the injection of PGE2. Unlike what we saw with the model of priming with inflammation (Fig.3), maternal separation led to a long-lasting decrease in *FKBP51* expression in both the hippocampus and the spinal cord (Hippocampus: Control vs Mat Sep: P<0.05 and Spinal Cord: Control vs Mat Sep: P<0.001) (Fig.4B). While no CpG sites passed our strict threshold for statistically different DNAm levels between control and maternal separation tissue, we identified two CpG sites with DNAm change >1% and with P trends <0.1: CpG at chr20:8,063,895 (DNAmΔ = 5.4%; FDR-corrected p=0.088) and CpG at chr20:8,096,364 (DNAmΔ = 10.5%; FDR-corrected p=0.071) (Fig.4C).

**Figure 4:**
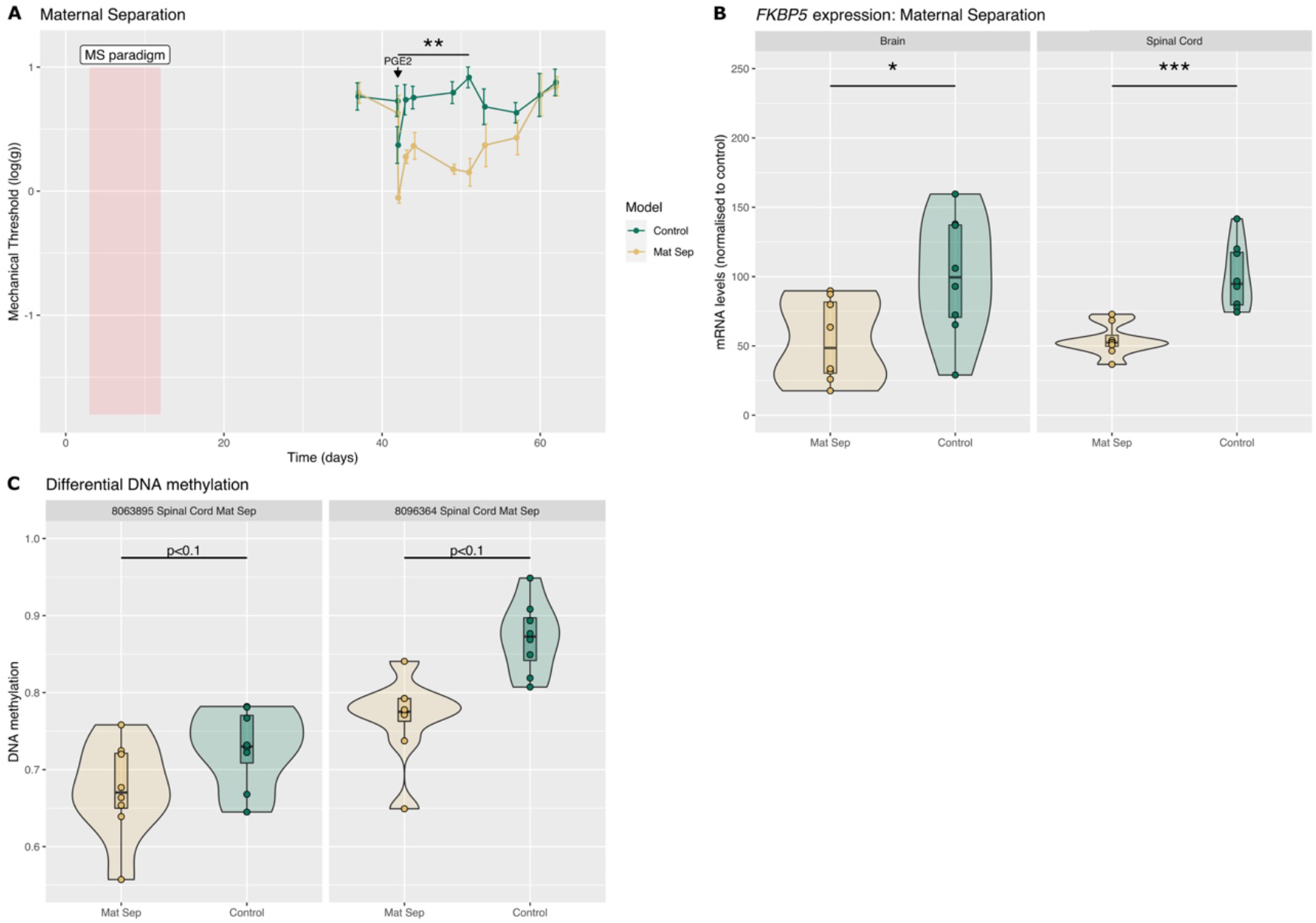
Maternal separation primes for hyper-responsiveness in later life and leads to a reduction in FKBP51 expression and *FKBP5* DNA methylation. **(A)** Maternal separation from P2 to P12 enhances mechanical hypersensitivity to PGE2 in adulthood. All animals received a s.c. injection of PGE2 (100ng/25ul) in the left hindpaw on Day 42 (also P42). N=8/8. 2-way ANOVA factor treatment, Day42 to Day49, F_1,14_=15.3, P<0.01, Control vs Mat Sep. **(B)** There was a significant decrease in FKBP51 mRNA in both the hippocampus and the spinal cord of MS rats at Day 40 (also P40). mRNA levels were quantified by RTqPCR. N=8/8. Data was normalized to Control. Hippocampus: Control vs Mat Sep: P<0.05 and Spinal Cord: Control vs Mat Sep: P<0.001 **C)** We identified two loci with different levels of DNAm between control and Maternal Separation with DNAm change >1% and with FRD-corrected P trends <0.1 (chr20:8,063,895; chr20:8,096,364).

These CpGs also both reside within a CpG shore (Table 2), and these shore regions have been observed to both positively and negatively correlate with gene expression, potentially through interaction with certain activating or repressive Transcription Factors [2]. In our case, there was a reduction in DNAm with reduced gene expression.

**Table 2:** Location of major DNA methylation difference between control and maternal separation tissue.

| Significant Cytosine (rn6) | Location | Distance to CGI (bp) | Nearest CGI | TFBS motifs ($\pm 10$bp) Transfac include | Mouse Syntenic Loci Regulatory Prediction (CNS) |
| --- | --- | --- | --- | --- | --- |
| chr20:8,063,895 | CGI shore | 1,741 | chr20:8,061,330-8,062,154 | TFCP2 | Quiescent/<br>Polycomb Repressed/ Poised Enhancer |
| chr20:8,096,364 | CGI shore | 935 | chr20:8,097,299-8,097,607 | EGR | Active Promoter/<br>Polycomb Repressed (Bivalent) |

Includes (Left to Right): Location of Cytosine (rn6 build); Location annotation; Distance to nearest CpG Island (CGI); Co-ordinates of nearest CGI; Highlighted enriched Transcription Factor Binding Site motifs in $\pm 10$ bp window centred on CpG via Transfac (TRAP analysis (24); and observed Chromatin Segmentation signature in the syntenic Mouse location (GRCm38/mm10, via LiftOver) in developmental central neural tissue (27)).

We next interrogated the DNA sequence for potential TFBS motifs as before and identified two potential motifs for the CpG at chr20:8,063,895 and eight potentials motifs for the CpG at chr20:8,096,364 (Supplementary Table S3). This included the motif for the Early Growth Response (EGR) transcription family, previous involved nociceptive signaling [33,36].

## Discussion

We have previously demonstrated that following noxious stimulation, *FKBP5* is rapidly upregulated in the superficial dorsal horn while *FKBP5* DNA methylation (DNAm) is reduced [28]. We also previously showed that FKBP51 drives persistent pain of neuropathic and inflammatory origin and that its inhibition during the maintenance stage of these chronic conditions reduces mechanical hypersensitivity [27]. We report here that SNI and CFA-induced inflammation induce long-lasting mechanical hypersensitivity of similar intensity but are not accompanied by a persistent upregulation of FKBP51. Crucially, the DNAm landscape of *FKBP5* was different between the two pain states, suggesting that FKBP5 promoter DNAm could indicate two distinct chronic pain pathways. We also found that the latent state of hypersensitivity induced by inflammation (or physical trauma) in early post-natal age was not accompanied by a change in *FKBP5* expression nor DNAm. However, there was a reduction of *FKBP5* DNAm together with a decrease in *FKBP5* expression following early life adversity (or emotional trauma), a paradigm that also primes for hyper-responsiveness to injury in adulthood. These results suggest that different mechanisms depending on the trauma may underlie the susceptibility to chronic pain.

Investigating *FKBP5* promoter DNAm landscape after ankle joint inflammation and neuropathic injury revealed differences in DNAm between the two pain states. Interestingly, the difference in DNAm at chr20:8,059,052 between CFA and SNI animals was significant in the hippocampus and also trended towards significance in the spinal cord. Indeed, the CpG at the position chr20:8,059,052 was the second CpG most likely to show a change in DNAm in spinal cord tissue. Moreover, our analysis showed a generalised difference of the same amplitude and direction at this CpG site in blood samples and there was a correlation between the DNAm level in the hippocampus and that in the blood and the spinal cord. These results suggested that changes in DNAm identified at this site may represent systemic changes occurring across multiple tissues (Supplementary Table S2).

*FKBP5* DNAm was greater in SNI than CFA animals and, while there was no difference in DNAm between the CFA and CFA Sham animals, both at CpG at chr20:8,059,052 and at 8,059,072 in the brain and the spinal cord, there was a trend for an increase between SNI and SNI Sham at these CpGs sites (Supplementary Table S2). The percentage methylation values for SNI sham, CFA sham and CFA animals were also extremely similar, suggesting that DNAm was likely to have been increased by the neuropathic injury in the SNI group. While more work is required to confirm these early observations, the significant correlation between the DNAm levels in the CNS and the blood suggests that the DNAm level at chr20:8,059,052 could act as a blood biomarker of neuropathic pain. Although our data suggest that there is no correlation between the DNAm level and the mechanical hypersensitivity associated with persistent pain, it may be that DNAm level at chr20:8,059,052 correlates with emotional behaviours such as anxiety and depression that often accompanies long-lasting pain states.

Further investigation will be needed to reveal the functional relevance of an increase in DNAm at this site, but it is of interest to note that chr20:8,059,052 may interact with the transcription factor E2F1 (Supplementary Table S3). Whilst the interaction between DNAm and transcription factors is complex, classically it is observed to inhibit TF binding, but can conversely also attract specific TFs [43]. E2F1 is particularly relevant in the context of neuropathic pain. It was found elevated in the blood of patients with spinal cord injury induced neuropathic pain [45] and in the hippocampus of hypersensitive mice following spinal cord injury [42]. Moreover, its ablation reduced mechanical sensitivity in this context [41]. The increase in DNAm observed in our study would therefore potentially prevent upregulation of FKBP51 induced by E2F1 binding at the chr20:8,059,052 CpG after nerve injury. Indeed, *FKBP5* mRNA was not found elevated in our CNS samples.

We also observed an increase in DNAm in SNI compared with CFA in the spinal cord at chr20:8,059,072. This CpG resides within a binding motif for the TF AP1 (Supplementary Table S3), which also has great relevance to nociceptive signaling. Indeed, AP1 binding activity rapidly increases in the spinal cord after noxious stimulation and it has recently been identified as a key gene associated with neuropathic pain in the spinal cord [26]. Similarly to the CpG at chr20:8,059,052, an increase in DNAm at this cytosine could prevent up-regulation of *FKBP5* induced by AP1 and would explain the lack of *FKBP5* upregulation observed in our samples.

Looking at the samples from animals that had been primed for hyper-responsiveness to inflammation in adulthood, we identified a decrease in DNAm at two other locations, chr20:8,063,895 and chr20:8,096,364 after maternal separation, but not after physical trauma. Interestingly, CpG at chr20:8,096,364 is likely to interact with the EGR transcription factor family. The most studied member of this family in the context of nociceptive signaling is the immediate early gene Egr1 (Zif268), which is rapidly up-regulated in the dorsal horn following noxious stimulation and, similarly to FKBP51, contributes to the maintenance of hypersensitive states [33,36]. A decrease in DNAm at chr20:8,096,364 could suggest that increase binding of EGR1 may occur following subsequent noxious stimulation, which in turn may lead to greater levels of FKBP51 and greater hypersensitivity, as seen in maternally separated animals, following noxious stimulation. It is important to note that the decrease in DNAm at chr20:8,063,895 and chr20:8,096,364 was accompanied by a significant decrease in *FKBP5* mRNA levels at baseline. However, the impact of this decrease in DNAm on *FKBP5* mRNA expression following noxious stimulation has not been explored here.

Overall, we found no correlation between the interrogated CpGs’ DNAm and expression levels of *FKBP5* mRNA, suggesting a complex regulation of gene expression for the gene *FKBP5*. Moreover, *FKBP5* DNAm levels at chr20:8,059,052 and 8,059,072 could not explain the long-lasting mechanical hypersensitivity seen in persistent pain states, as SNI and CFA animals had similar degree of mechanical hypersensitivity but different DNAm profiles at these loci. Finally, *FKBP5* mRNA was not up-regulated 30 days after CFA and SNI, suggesting an intricate mechanism of regulation of persistent pain by FKBP51, as pharmacological inhibition of this protein in the maintenance phase of the pain states still reduces mechanical hypersensitivity [27]. While our previous work had highlighted FKBP51 as a new target for the treatment of chronic pain, this study is the first to propose *FKBP5* promoter DNAm landscape as a pathological indicator and potential biomarker of neuropathic pain and of the susceptibility to chronic pain.

## Abbreviations

DNAm: DNA methylation

## Acknowledgments

This work was funded by a research grant from Mundipharma Research Limited (MRL) to SG.

The authors report no conflict of interests.

## Data availability

Data is available for sharing. Contact the corresponding author.

