## Supplementary Data for "Using the DNA methylation profile of the stress driver gene *FKBP5* for chronic pain diagnosis"

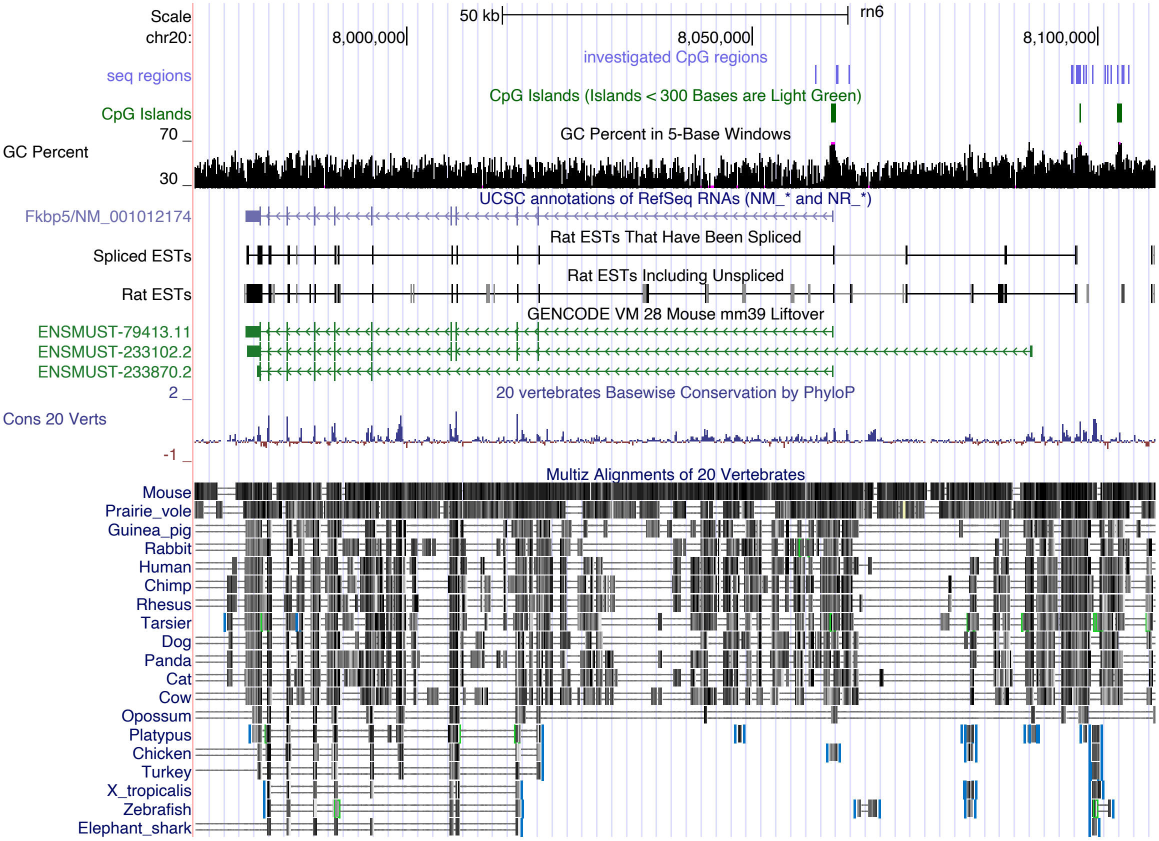


**Fig. S1. Location of sequencing probes of 3 5'CpG Island regions and plausible promoter regions of *Fkbp5* gene displayed in UCSC Browser.** Layers from top: Scale in Rat (rn6) build; Genomic location chr20:7,969,338-8,108,322; Location of Targeted Sequencing regions for DNAm analysis (purple); Location of CpG Islands (green); GC Percentage indicating increase at CpG Island locations; Location of Fkbp5 from UCSC, Rat Spliced and unspliced ESTs; Mouse mm39 Gencode (vm28) LiftOver; Conservation across 20 Vertebrates (PhyloP) and as individual Multiz Alignments.

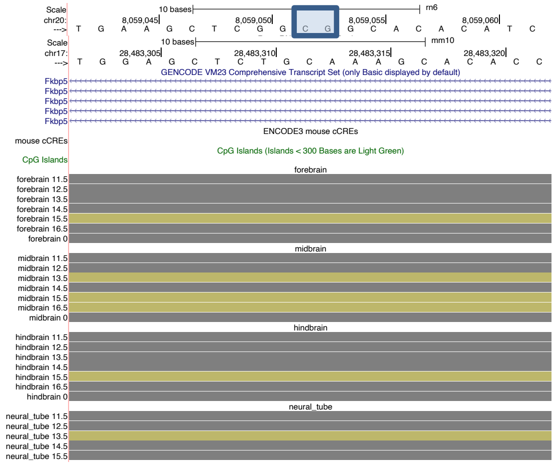


Chr20:8059052

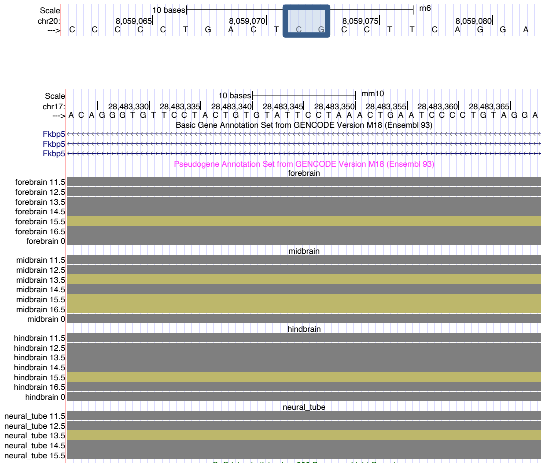


Chr20:8059072

| | | |
|---|---|---|
| 1 | Tss | Active TSS |
| 2 | TssFlnk | Flanking active TSS |
| 3 | Tx | Transcription |
| 4 | TxWk | Weak transcription |
| 5 | EnhG | Enhancer in gene |
| 6 | Enh | Enhancer |
| 7 | EnhLo | Weak enhancer |
| 8 | EnhPois | Poised enhancer |
| 9 | EnhPr | Primed enhancer |
| 10 | TssBiv | Bivalent TSS |
| 11 | ReprPC | Repressed by Polycomb |
| 12 | ReprPCW | Repressed by Polycomb (wk) |
| 13 | QuiesG | Quiescent gene |
| 14 | Quies | Quiescent |
| 15 | Quies2 | Quiescent |
| 16 | Quies3 | Quiescent |
| 17 | Quies4 | Quiescent |
| 18 | Het | Heterochromatin |

**Fig. S2.: Chromatin Segmentation in Syntenic Mouse Regions.** Top to Bottom: Sequence location in Rat (rn6); Sequence location in Mouse (GRCm38/mm10); Gencode Transcript Information (VM23); mouse CREs; CpG islands; Chromatin Segmentation in developing Forebrain, Midbrain, Hindbrain, and Neural Tube. Legend colour code for 18-state Segmentation.

**Table S1. Investigated CpG regions (Rat assembly rn6)**

Type or paste caption here. Create a page break and paste in the Table above the caption.

| chromosome | start | stop | CpGs covered |
|---|---|---|---|
| chr20 | 8059015 | 8059131 | 4 |
| chr20 | 8062075 | 8062179 | 9 |
| chr20 | 8062233 | 8062336 | 3 |
| chr20 | 8063885 | 8063995 | 5 |
| chr20 | 8096103 | 8096195 | 2 |
| chr20 | 8096316 | 8096391 | 2 |
| chr20 | 8096800 | 8096880 | 5 |
| chr20 | 8096916 | 8096993 | 5 |
| chr20 | 8097096 | 8097190 | 2 |
| chr20 | 8097291 | 8097389 | 12 |
| chr20 | 8097867 | 8097945 | 2 |
| chr20 | 8098183 | 8098283 | 3 |
| chr20 | 8099134 | 8099255 | 3 |
| chr20 | 8100968 | 8101069 | 2 |
| chr20 | 8101327 | 8101429 | 2 |
| chr20 | 8101818 | 8101915 | 2 |
| chr20 | 8102706 | 8102783 | 4 |
| chr20 | 8102783 | 8102870 | 8 |
| chr20 | 8103416 | 8103525 | 5 |
| chr20 | 8103670 | 8103774 | 7 |
| chr20 | 8104402 | 8104523 | 3 |

**Table S2. DNA methylation levels in injured and control animals at CpG sites chr20: 8,059,052 and chr20: 8,059,072.**

| CpG/tissue | CFA | CFA Sham | SNI | SNI Sham | SNI -CFA | SNI-SNI sham | CFA-CFA sham |
|---|---|---|---|---|---|---|---|
| **8,059,072/spinal cord** | 86.9 ± 1.3 % | 88.8 ± 2.2 % | 97.3 ± 1.1 % | 88.8 ± 4.2 % | 10.5 % | 8.5% | -1.9% |
| **8,059,052/spinal cord** | 82.1 ± 2.9 % | 85.6 ± 2.6 % | 94.0 ± 2.6 % | 86.6 ± 5.2 % | 11.9 % | 7.4% | -3.5% |
| **8,059,052/hippocampus** | 83.5 ± 0.9 % | 84.0 ± 1.4 % | 94.6 ± 1.3 % | 91.5 ± 5.3 % | 11.2% | 3.2% | -0.6% |
| **8,059,052/blood** | 80.4 ± 1.7 % | 81.6 ± 2.0 % | 88.9 ± 1.3 % | 83.1 ± 2.0 % | 8.5% | 5.8% | -1.2% |

**Table S3. Transfac Motif prediction via TRAP.**

| CpG | Rank | P-value | Matrix ID | Matrix name |
|---|---|---|---|---|
| 8,059,052 | 1 | 0.0218 | M00428 | V$E2F1_Q3 |
| 8,059,052 | 2 | 0.0409 | M01598 | V$ZBED6_01 |
| 8,059,052 | 3 | 0.0448 | M00431 | V$E2F1_Q6 |
| 8,059,072 | 1 | 0.0189 | M00495 | V$BACH1_01 |
| 8,059,072 | 2 | 0.0309 | M00174 | V$AP1_Q6 |
| 8,059,072 | 3 | 0.0496 | M00199 | V$AP1_C |
| 8,063,895 | 1 | 0.0128 | M00720 | V$CACBINDINGPROTEIN_Q6 |
| 8,063,895 | 2 | 0.0344 | M00072 | V$CP2_01 |
| 8,096,364 | 1 | 0.0171 | M00776 | V$SREBP_Q3 |
| 8,096,364 | 2 | 0.0188 | M00497 | V$STAT3_02 |
| 8,096,364 | 3 | 0.023 | M00690 | V$AP3_Q6 |
| 8,096,364 | 4 | 0.0249 | M00494 | V$STAT6_01 |
| 8,096,364 | 5 | 0.0258 | M00979 | V$PAX6_Q2 |
| 8,096,364 | 6 | 0.0267 | M00807 | V$EGR_Q6 |
| 8,096,364 | 7 | 0.0283 | M00246 | V$EGR2_01 |
| 8,096,364 | 8 | 0.0496 | M00378 | V$PAX4_03 |
